# Minimal information for Chemosensitivity assays (MICHA): A next-generation pipeline to enable the FAIRification of drug screening experiments

**DOI:** 10.1101/2020.12.03.409409

**Authors:** Ziaurrehman Tanoli, Jehad Aldahdooh, Farhan Alam, Yinyin Wang, Umair Seemab, Maddalena Fratelli, Petr Pavlis, Marian Hajduch, Florence Bietrix, Philip Gribbon, Andrea Zaliani, Matthew D. Hall, Min Shen, Kyle Brimacombe, Evgeny Kulesskiy, Jani Saarela, Krister Wennerberg, Markus Vähä-Koskela, Jing Tang

**Affiliations:** Research Program in Systems Oncology, Faculty of medicine, University of Helsinki, Finland; Istituto di Ricerche Farmacologiche Mario Negri IRCCS, Italy; Institute of Molecular and Translational Medicine, Czech; European infrastructure for translational medicine; Fraunhofer Institute for Translational Medicine and Pharmacology, Hamburg, Germany; National Center for Advancing Translational Sciences, U.S.A; Institute for Molecular Medicine Finland, University of Helsinki, Finland; Biotech Research & Innovation Centre (BRIC), University of Copenhagen, Denmark

**Keywords:** Drug discovery, drug sensitivity assays, data integration tools, FAIR research data

## Abstract

Chemosensitivity assays are commonly used for preclinical drug discovery and clinical trial optimization. However, data from independent assays are often discordant, largely attributed to uncharacterized variation in the experimental materials and protocols. We report here the launching of MICHA (Minimal Information for Chemosensitivity Assays), accessed via https://micha-protocol.org. Distinguished from existing efforts that are often lacking support from data integration tools, MICHA can automatically extract publicly available information to facilitate the assay annotation including: 1) compounds, 2) samples, 3) reagents, and 4) data processing methods. For example, MICHA provides an integrative web server and database to obtain compound annotation including chemical structures, targets, and disease indications. In addition, the annotation of cell line samples, assay protocols and literature references can be greatly eased by retrieving manually curated catalogues. Once the annotation is complete, MICHA can export a report that conforms to the FAIR principle (Findable, Accessible, Interoperable and Reusable) of drug screening studies. To consolidate the utility of MICHA, we provide *FAIRified* protocols from five major cancer drug screening studies, as well as six recently conducted COVID-19 studies. With the MICHA webserver and database, we envisage a wider adoption of a community-driven effort to improve the open access of drug sensitivity assays.

## INTRODUCTION

Drug sensitivity or chemosensitivity assay is an important tool to measure cellular response to drug perturbation, which has been increasingly used for preclinical drug discovery and clinical trial optimization. However, poor inter- and intra-laboratory reproducibility has been reported when comparing batches that differ at assay materials and methods [1–3]. Central to improving the data reproducibility is the standardization of assay ontologies, with which the research community may annotate minimal information (MI) about critical components of a typical drug sensitivity experiment. An existing implementation is MIABE (minimum information about a bioactive entity), which provides guidelines for a wide range of bioactivity assay types including cellular assays, whole organism and pharmacokinetic studies, but it has not been tailored for drug sensitivity assays [4]. Furthermore, like many other similar MI efforts in omics assays, MIABE provides limited data integration tools to enable its implementation coherently.

Several databases store the actual drug sensitivity data points such as CCLE[5], GDSC [6] and CTRP[7], but there is a lack of annotation for these assay protocols, making it challenging to evaluate the methodological differences in the experiments. The solution we present here, MICHA (minimal information for chemosensitivity assays), provides an integrative pipeline to facilitate the annotation of four major components of any drug sensitivity assay, including 1) compounds, 2) samples, 3) reagents, and 4) data processing references. Using the platform of MICHA, we aim to increase acceptance and adoption of the principles of FAIR (Findable, Accessible, Interoperable and Reusable), by making the assay annotation as smoothly as possible with data integration tools and databases. Furthermore, with the help of MICHA, we catalogue the major drug sensitivity screening protocols in cancer and COVID-19 that may help users compare the existing experiments as well as inform the design of new experiments.

## MATERIALS AND METHODS

### Workflow

Using MICHA, users can upload their chemicals, samples, and experimental design information (**Figure 1**). To start, users need to upload the names and InChiKeys of the compound, after which MICHA will automatically extract primary and secondary target information, physio-chemical properties, and disease indications for these compounds. This information will help users understand the mechanisms of action for their compounds. After obtaining the drug target profiles along with other annotations, users may continue filling in the other experimental details, such as sample (cell lines or patient-derived samples) information and assay conditions. For cell lines, only the names of the cell lines are required, as the other information will be retrieved automatically from internal databases. For annotating assay protocols, we derived a consensus on the minimal information that is needed, including assay format, detection technology, end point mode of action, experimental medium, plate type, cell density, time for treatment, dilution fold, vehicle of compound, dispensation method and volume per well. The definition of these terminologies is explained in **Supplementary File 1** as well as in the ‘**Glossary’** tab at the MICHA website. Furthermore, users are directed to a web form to report data processing information, including minimum and maximum concentrations of the compounds, publication references and drug response metric names such as IC_50_ or AUC. Finally, a tabular report can be generated according to the user’s input combined with the augmented annotation retrieved from public resources. In addition, users may browse annotated protocols including five major cancer and six recent COVID-19 drug sensitivity screening studies. In future, we plan to annotate more protocols according to the workflow as proposed by MICHA.

**Figure 1:**
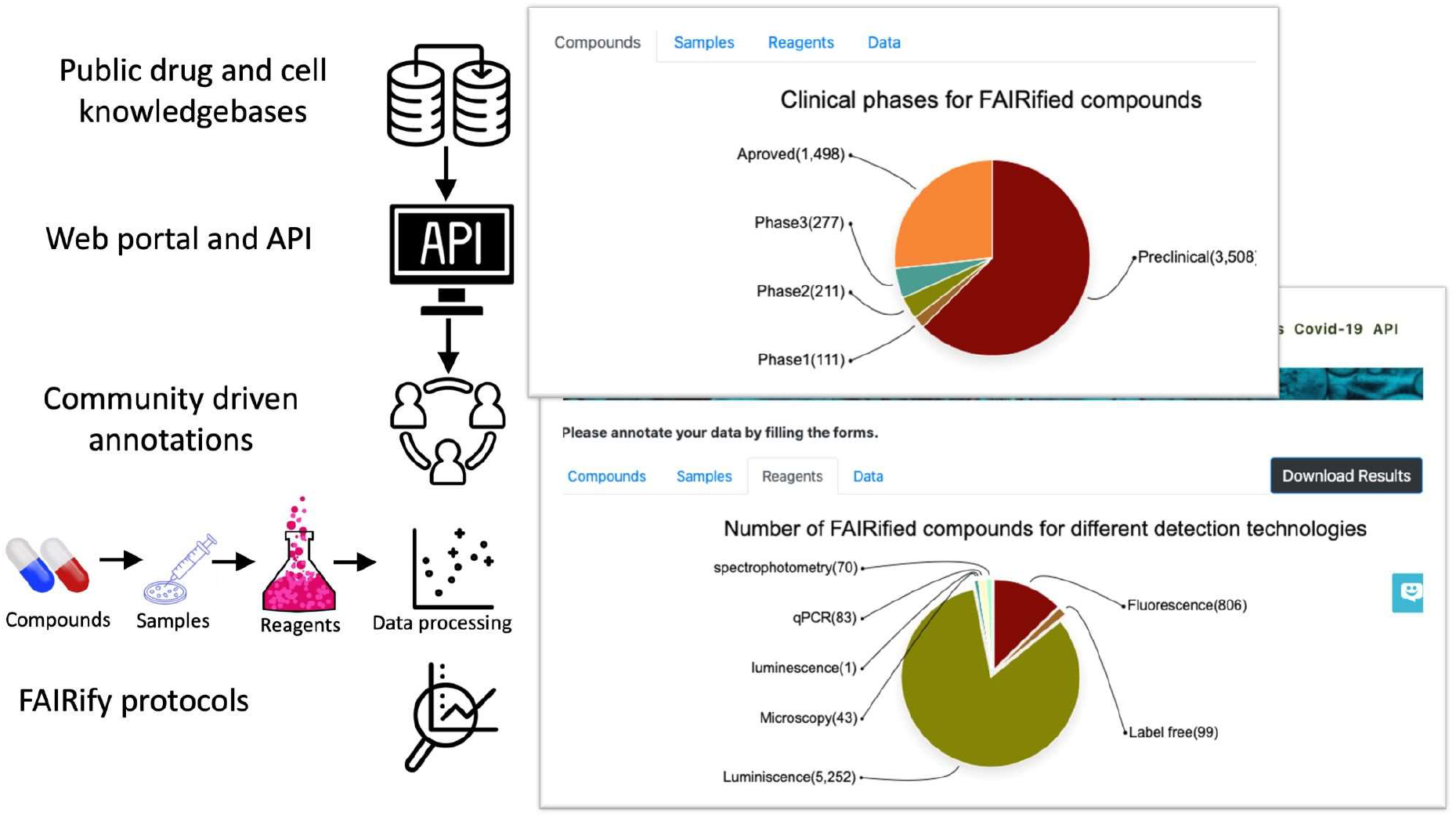
User interface and workflow of MICHA. Users start with compound annotation by uploading a list of compounds with names and standard InChiKeys. MICHA will return the pharmacological and physio-chemical properties of the compounds via an integrative webserver and database, available under the ‘Compounds’ tab. Users then may click on Samples, Reagents or Data processing tabs to annotate their drug screening protocols. Auto suggestions are provided to avoid spelling mistakes or terminology conflicts. Finally, users can download the summary reports containing input data as well as integrated information provided by MICHA. Pie charts under Compounds, Samples, Reagents and Data tab show the statistics for existing FAIRified protocols.

### Data integration tools

Three types of datasets are retrievable via data integration tools in MICHA:

#### 1) FAIRified protocols

A prime objective for MICHA is to provide a pipeline for the FAIRification of drug sensitivity assays, such that these established protocols can be well documented with enhanced visibility to the research community. To initiate such an effort, we have FAIRified drug screening protocols from major cancer studies including GDSC (345 compounds and 987 cell lines) [7], CCLE (24 compounds and 504 cell lines) [5] and CTRPv2 (203 compounds and 242 cell lines) [6]. Furthermore, we have provided drug sensitivity screening protocols extracted from six recent COVID-19 antiviral studies (5,525 compounds and 2 cell lines) [8–13]. On the other hand, we have provided an example of protocols established at the research institution level (528 compounds and 4 cell lines utilized at the high throughput drug-screening unit at the Institute for Molecular Medicine Finland, University of Helsinki). These FAIRified protocols can be freely obtained at http://micha-protocol.org/protocols/. With more protocols annotated via MICHA, the research community shall be better informed on the variations on the experimental condition across different studies and institutions. Table 1 represents an overview of FAIRified protocols by MICHA.

**Table 1:** FAIRified protocols by MICHA

| Protocol name | Type | Compounds | Cell lines | Min concentration (nM) | Max concentration (nM) | Metric | Reference |
| --- | --- | --- | --- | --- | --- | --- | --- |
| **GDSC** | Cancer | 345 | 987 | 0.03 | 4,000,000 | IC50 | [6] |
| **CCLE** | Cancer | 24 | 504 | 2.5 | 8,000 | IC50, EC50 | [5] |
| **CTRPv2** | Cancer | 203 | 242 | 0.56 | 592,000 | IC50 | [7] |
| **FIMM** | Cancer | 528 | 4 | 0.0 | 1,000,000 | DSS | [14] |
| **Mario Negri** | Cancer | 1 | 16 | 0.0 | 10,000 | AUC | [15] |
| **NCATS** | COVID-19 | 5,108 | 1 | 0.0 | 120,000 | AC50 | [12] |
| **Ellinger et al.,** | COVID-19 | 103 | 1 | 20 | 20,000 | IC50, CC50 | [13] |
| **Gorden et al.,** | COVID-19 | 73 | 1 | 1 | 10,000 | IC50 | [9] |
| **Jeon et al.,** | COVID-19 | 43 | 1 | 50 | 50,000 | IC50 | [11] |
| **Touret et al.,** | COVID-19 | 83 | 1 | 600 | 40,000 | EC50 | [10] |
| **Wetson et al.,** | COVID-19 | 43 | 1 | 50 | 5,000 | IC50 | [8] |

#### 2) Drug-target data

Compound target profiles are integrated from the most comprehensive drug target databases including DTC [16][17], BindingDB [18], ChEMBL [19], GtopDB [20], DGiDB [21] and DrugBank [22]. The first four databases (DTC, BindingDB, ChEMBL and GtopDB) contain quantitative bioactivity data whereas DGiDB and DrugBank contain unary drug target information. We have focused on the primary and secondary targets of a compound, defined as those displaying binding affinities <= 1,000 nM from the bioactivity databases, or those that are recorded in the unary databases. We have integrated drug targets for 277K chemicals from DTC, 513K from ChEMBL, 258K from Binding DB, 4.8K from GtopDB, 7.6K from DGiDB and 6.8K from DrugBank. Furthermore, we have merged overlapping targets across these databases to avoid duplications, resulting in high-quality target profiles for >800K chemicals. Such a data integration provides one of the most comprehensive compounds collection along with their potent primary and secondary targets. All the drug-target information can be retrieved by the MICHA annotation pipeline or programmatically using the API (application programming interface).

#### 3) Compound properties, cell line and assay information

Compound physicochemical properties and structures for 1.9 million compounds are obtained from the ChEMBL database. Furthermore, we have integrated disease indications and clinical phase information for 3600 pre-clinical and clinical drugs from the DTC database. This information together with the drug target profiles will be retrieved for the user-uploaded compound list. When users annotate the cell lines, the majority of cell line information can be retrieved automatically from Cellosaurus [23], which is a comprehensive knowledge database on cell lines. For assay annotation, commonly used techniques will be provided for users to choose from to ease the burden of manual editing.

## ADDED VALUES BY MICHA

### Comprehensive drug-target profiles

For annotating the mechanisms of action of compounds, MICHA integrates the most up-to-date potent drug target profiles from various databases, which harbor drug-target data at different levels, ranging from quantitative bioactivity values to unary drug target hits. For instance, DrugBank, GtopDB and DGiDB are mainly focused on approved compounds with putative target information, while ChEMBL, BindingDB and DTC include bioactivity values for more versatile investigational and pre-clinical chemicals. In MICHA, we have tried to improve target coverage across the druggable genome by integrating non-overlapping data points from the latest releases of these databases. As shown in **Figure 2**, the average number of targets for 2,993 approved drugs (active ingredients including salts) in MICHA is 7.33, as compared to that from ChEMBL (5.5), DGiDB (4.71), DrugBank (3.56), BindingDB (2.74) and GtopDB (0.96). Similarly, for 1,992 investigational compounds (defined as those in clinical trials), the average number of targets per compound is higher in MICHA as compared to other databases. Compound target profiles in MICHA will be updated upon the new release of the corresponding databases. Secondly, the API for MICHA provides comprehensive target profiles for a compound within fraction of a second by inputting its standard InChiKey, available at: https://api.micha-protocol.org. We believe that the API for drug target information will further boost the usability of MICHA by programmatically integrating compound target profiles with other related tools, and shall open new applications for drug discovery researchers for training their compound-target machine learning models [24–26] as well as providing more insights on the network modeling of mechanisms of action.

**Figure 2:**
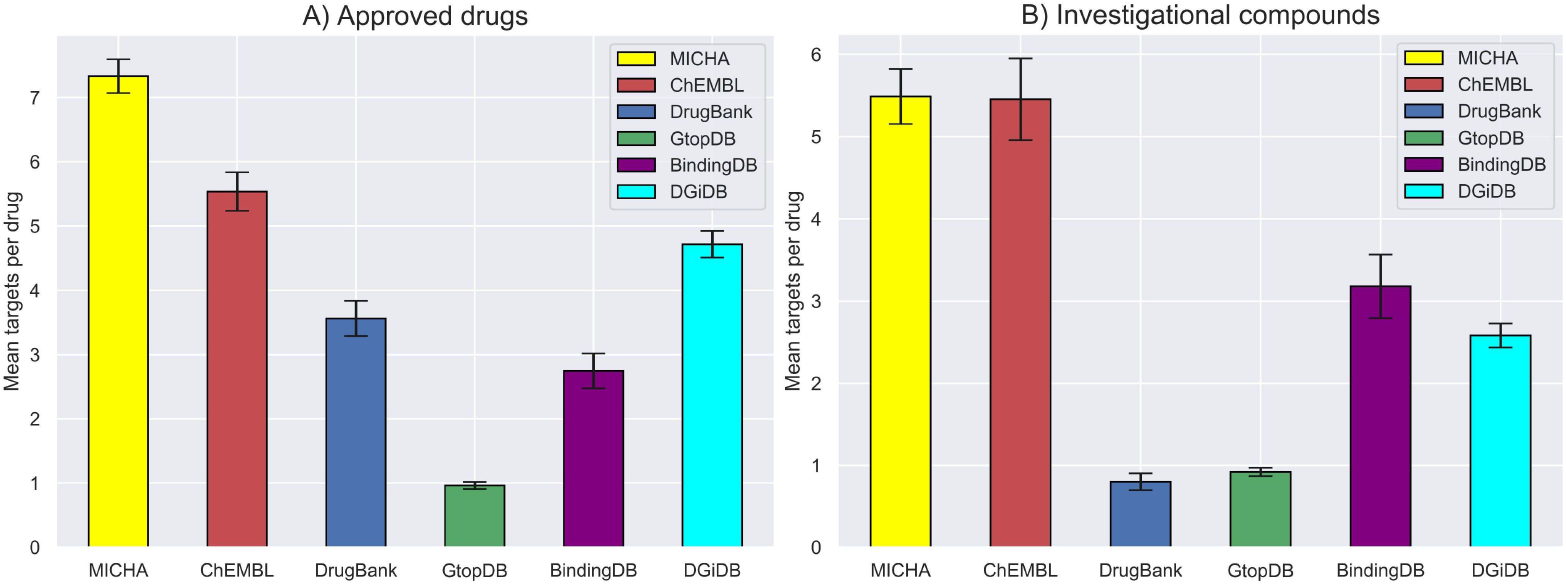
Average number of targets for approved and investigational compounds in multiple databases.

### FAIRification of drug sensitivity screening protocols

We have FAIRified the screen protocols for three major cancer drug studies including CCLE, GDSC and CTRPv2, available at https://micha-protocol.org/protocols. These drug screening studies share similar objectives of linking genetic features of cancer cell lines to small-molecule sensitivity to accelerate drug discovery. Note that MICHA focuses on the annotation of drug screening protocols while the actual data points are available in their corresponding databases. Here we report the comparison of the major components in the assay protocols (**Table 2**).

**Table 2:** Comparison between protocols of CCLE, GDSC and CTRPv2. NA indicates that the information is unavailable.

|  | CCLE | GDSC | CTRPv2 |
| --- | --- | --- | --- |
| Plate types | 1536 | 384, 96 | 384-Opaque white |
| *Experimental media* | NA | DMEM, RPMI | ALPHAMEM, DMEM, DMEMF, EMEM, HAMSF, IMDM, L15, McCoy5A, MCDB, MEM, RPMI, Waymouth, WilliamsE |
| *Detection technology* | Luminescence | Fluorescence | Luminescence |
| *Cell density* | 250 | NA | 500, 1000, 2000, 3000, 5000, 10000 |
| *Treatment time (h)* | 72-84 | 72 | 72 |
| *Analysis metrics* | IC<sub>50</sub>, EC<sub>50</sub> | IC<sub>50</sub> | IC<sub>50</sub> |
| *Compounds* | 24 | 345 | 203 |
| *Number of cell lines* | 504 | 987 | 242 |

Both GDSC and CTRPv2 used common experimental plate type i.e., 384-well plates, whereas CCLE compounds were tested on 1536-well plates. In GDSC, two different experimental media, including DMEM and RPMI were tested for the 987 cancer cell lines, whereas the CTRPv2 cell lines were tested for many more different media as listed in **Table 2**. In contrast, we could not find experimental medium information for CCLE. On the other hand, both CCLE and CTRPv2 have used Cell-Titer-Glo (Promega), a luminescence-based assay to measure the levels of ATP as a surrogate for cell viability, whereas GDSC has used based nucleic acid staining syto60 (Invitrogen) for adherent cells and resazurin (Sigma) for suspension cells. All the three screening studies have used at least 72 hours of treatment, after which the IC_50_ or EC_50_ concentrations were determined from the dose response curves.

**Figure 3** shows the overlapping chemicals and cell lines tested across CCLE, GDSC and CTRPv2 studies, after excluding those chemicals for which proper chemical names or identifiers were missing to assure high quality data in MICHA. Only two chemicals are shared across the three studies including selumetinib and tanespimycin (**Figure 3A**). Selumetinib (AZD6244) is a MEK (kinase) inhibitor used for treating neurofibromatosis type I in children [27]; whereas tanespimycin is a Hsp90 inhibitor [28] that has been studied for the treatment of leukemia or solid tumors, especially kidney tumors. In contrast, more overlap was found for the cell lines, with 112 cell lines in common across the three studies (**Figure 3B**).

**Figure 3:**
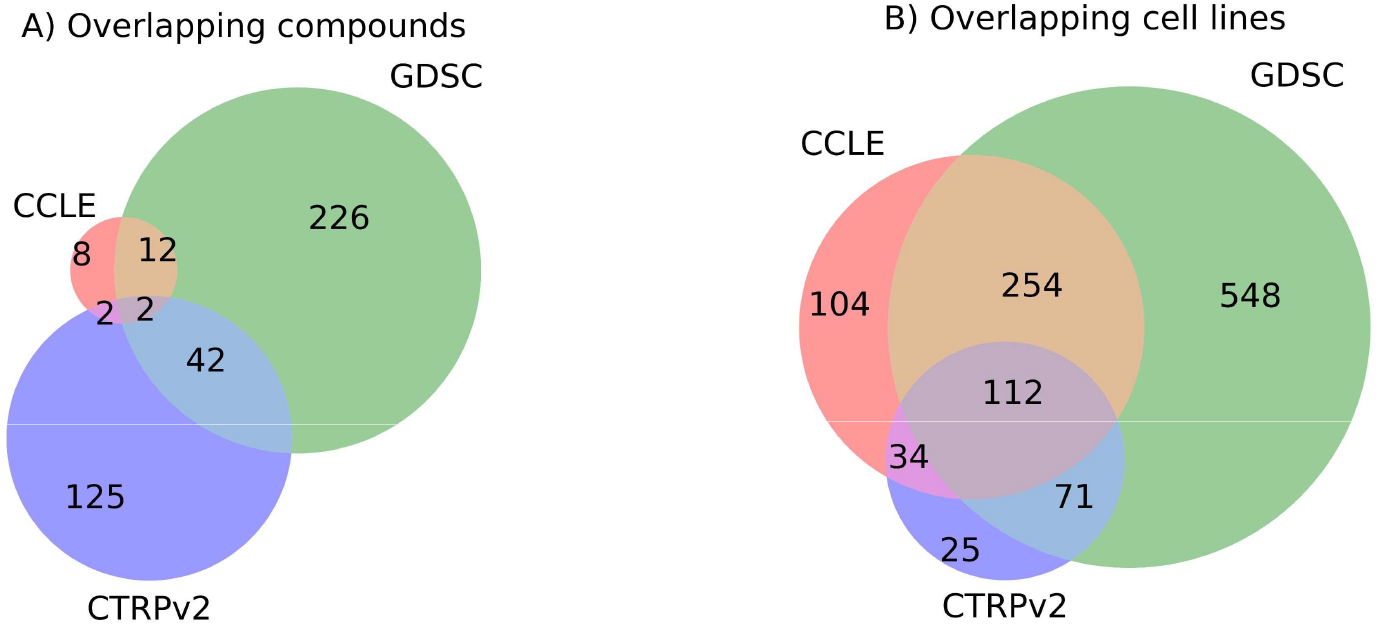
Overlapping compounds and cell lines between CCLE, GDSC and CTRPv2.

MICHA has not only FAIRified cancer related drug screening studies but also annotated six recent studies on COVID-19, a virus that causes ongoing pandemic with limited drug treatment options. From these studies, we have identified 5,318 chemicals tested across two cell lines including Vero E6 and Caco-2. The annotations of these compounds, cell lines, as well as the experimental information and data analysis methods can be easily retrieved at http://micha-protocol.org/covid19.

With the help of MICHA platform, all these COVID-19 and cancer related drug screening protocols are freely accessible to the users (Findable). In total, **Figure 4A** shows the clinical phases of compounds FAIRified by MICHA, whereas **Figure 4B** shows the distribution of FAIRified cell lines from different tissue types. These statistics show a broad coverage of cell lines and compounds. We believe that with the FAIRification of more protocols, MICHA has the potential to become a standard workflow for annotating and cataloguing chemosensitivity experiments. Moreover, these protocols can be accessed programmatically using MICHA API available at: https://api.micha-protocol.org/, which makes it possible for *In silico* models to programmatically access MICHA to obtain compound information such as protein targets and physicochemical properties and use this information for novel drug target predictions (Interoperable). The MICHA drug screening protocols can be considered as a reference for the experimental design of future drug screening studies, as well as serve as a source of information to evaluate the experimental reproducibility (Reusable). Moreover, the MICHA source code is available at GitHub for reuse of the application towards a tailoring need for individual drug screening centers (https://github.com/JehadAldahdooh/MICHA). MICHA is also indexed at https://fairsharing.org/ to be accepted as a potential tool for chemosensitivity data FAIRification.

**Figure 4:**
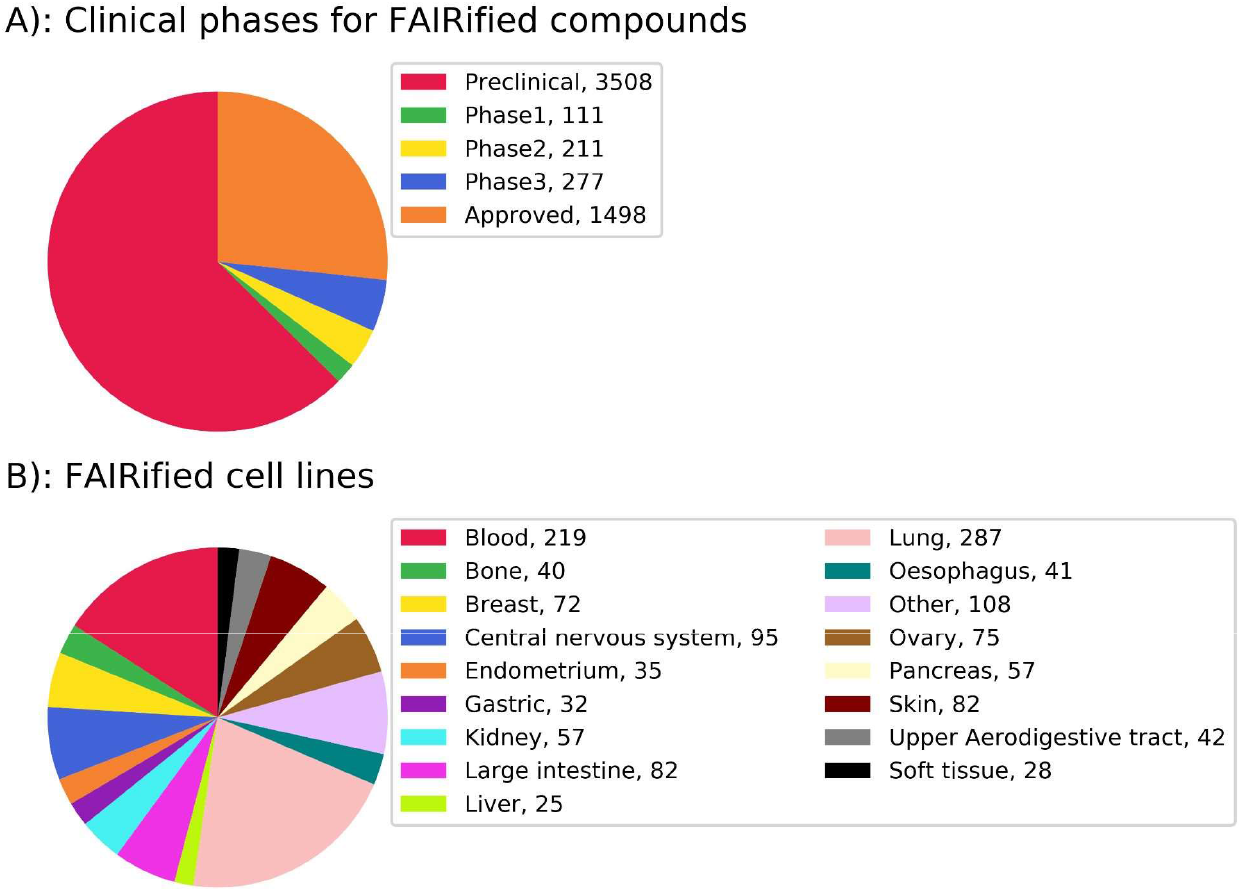
A) Clinical phase of compounds in FAIRified protocols. B) Tissue types for cell lines in FAIRified protocols.

## CONCLUSION

Chemosensitivity assay screening has been increasingly utilized for preclinical drug discovery and clinical trial optimization. However, chemosensitivity assays often lack sufficient annotation to make the data FAIR, which has become a limiting factor for supporting its clinical translation. To improve the assay annotation, web portals that facilitate information retrieval from different assay components are critically needed. To address this issue, we have recently launched MICHA (Minimal information for Chemosensitivity assays) as a web server for the annotation of chemosensitivity screens that covers critical information including 1) compounds 2) samples 3) reagent protocols and finally, 4) data processing methods.

Comprehensive compound-target profiles were deposited to the MICHA database for more than 800K compounds. These high-quality pharmacological data shall help improve the understanding of the mechanisms of action not only for approved drugs but also for investigational and preclinical compounds. Furthermore, the target profiles at the druggable genome scale provide more information on the polypharmacological effects, which might lead to new opportunities for drug repositioning [29]. To facilitate the data retrieval, the API in MICHA is highly optimized such that it can return target profiles for thousands of chemicals within seconds.

With the MICHA web portal, we have FAIRified major drug sensitivity screening protocols from five cancer studies and six recent COVID-19 studies, serving as the first instances of the catalogue. Comparing these deeply curated assay protocols should allow a more systematic analysis of data reproducibility. With the FAIR-compliant data resources and tools to deliver content standards and ontology services, MICHA will ensure the characterization of critical assay components, allow the FAIRification and cataloguing of drug sensitivity studies, and support the downstream analysis towards clinical translation. We invite the drug discovery community to use MICHA for annotating their drug sensitivity assays to improve the knowledge sharing, which shall ultimately lead to a bigger impact in translational medicine.

## Key points

- We have proposed a novel workflow called MICHA (https://micha-protocol.org) for the FAIRification of drug sensitivity screening protocols.
- MICHA provides an integrated platform to obtain drug screening assay annotations, drug target profiles, and other pharmacological information in an easy and fast manner.
- MICHA has FAIRified drug screening protocols related to cancer and COVID-19, which are made comparable and informative for designing new experiments.

## DATA AVAILABILITY STATEMENT

FAIRified protocols by MICHA are freely accessible using MICHA API at https://api.micha-protocol.org/.

Additionally, other related datasets can be downloaded using MICHA GUI.

## ACKNOWLEDGEMENT

We thank the CSC-IT Center for Science Finland for providing database storage and computing resources.

**Ziaurrehman Tanoli** is a senior researcher at the University of Helsinki. His research is mostly focused on computational drug repurposing. He is also developing bioinformatics tools for drug target interactions.

**Jehad Aldahdooh** is a PhD student at the University of Helsinki. He is developing text mining applications for drug target interactions.

**Farhan Alam** is a PhD student at the University of Helsinki.

**Yinyin Wang** is a PhD student at the University of Helsinki. She is developing methods for network pharmacology modeling for herbal medicine.

**Umair Seemab** is a PhD student at the University of Helsinki. He is working on NGS and single cell sequencing.

**Maddalena Fratelli** is the head of phamacogenomics unit at the Mario Negri Institute for pharmacological research. She is working on genomic and transcriptomic systems for the study of drug action and resistance.

**Petr Pavlis** is a software engineer and IT head at Institute of Molecular and Translational Medicine, Palacky University, Czech republic, focusing on informatics background in preclinical and clinical studies.

**Marian Hajduch** is the founding director of the Institute of Molecular and Translational Medicine. His research interests are in molecular and translational medicine.

**Florence Bietrix** is the head of operations at EATRIS. Her research interests are to develop new therapeutic targets for atherosclerosis and nonalcoholic steatohepatitis.

**Philip Gribbon** is the head of discovery research at the Fraunhofer Institute for Translational Medicine and Pharmacology.

**Andrea Zaliani** is a senior bioinformatics scientist at the Fraunhofer Institute for Translational Medicine and Pharmacology. He has expertise in pharmaceutical research and development.

**Matthew D. Hall** is a biology group leader at the National Center for Advancing Translational Sciences. He optimizes both biochemical and cell-based assays for automated, small molecule, high-throughput screening in collaboration with NIH.

**Min Shen** has extensive experience in cheminformatics and computational chemistry through her work at the National Center for Advancing Translational Sciences.

**Kyle Brimacombe** is a biochemist at the National Center for Advancing Translational Sciences.

**Evgeny Kulesskiy** is a senior researcher at the Institute for Molecular Medicine Finland. He is working on drug sensitivity and resistance testing.

**Jani Saarela** is the operational manager of the High Throughput Biomedicine unit at the Institute for Molecular Medicine Finland. He is working on drug sensitivity and resistance testing.

**Krister Wennerberg** is a professor at the Biotech Research & Innovation Centre. He is doing research on chemical systems biology applications.

**Markus Vähä-Koskela** is a senior researcher at the Institute for Molecular Medicine Finland. He is doing research in areas such as onco-immunology, immunotherapy, and translational cancer medicine.

**Jing Tang** is an assistant professor at the University of Helsinki. He is working on mathematical, statistical and informatics tools to tackle biomedical questions.

